# Thalamocortical network connectivity controls spatiotemporal dynamics of cortical and thalamic traveling waves

**DOI:** 10.1101/780239

**Authors:** Sayak Bhattacharya, Matthieu B. Le Cauchois, Pablo A. Iglesias, Zhe S. Chen

**Affiliations:** Department of Electrical and Computer Engineering, Whiting School of Engineering, Johns Hopkins University, Baltimore, MD 21218; Department of Mechanical Engineering, Whiting School of Engineering, Johns Hopkins University, Baltimore, MD 21218; Department of Psychiatry, Department of Neuroscience & Physiology, Neuroscience Institute, New York University School of Medicine, New York, NY 10016

## Abstract

Propagation of neural activity in spatially structured neuronal networks has been observed in awake, anesthetized and sleeping brains. However, it remains unclear how traveling waves are coordinated temporally across recurrently connected brain structures, and how network connectivity affects spatiotemporal neural activity. Here we develop a computational model of a two-dimensional thalamocortical network that enables us to investigate traveling wave characteristics in space-time. We show that thalamocortical and intracortical network connectivity, excitation/inhibition balance, thalamocortical/corticothalamic delay can independently or jointly change the spatiotemporal patterns (radial, planar and rotating waves) and characteristics (speed, direction and frequency) of cortical and thalamic traveling waves. Simulations of our model further predict that increased thalamic inhibition induces slower cortical wave frequency, and enhanced cortical excitation increases cortical wave speed and oscillation frequencies. Overall, the model study provides not only theoretical insight into the basis for spatiotemporal wave patterns, but also experimental predictions that potentially control these dynamics.

**Author Summary:** Cognition or sensorimotor control requires the coordination of neural activity across widespread brain circuits. Propagating waves of oscillatory neural activities have been observed at both macroscopic and mesoscopic levels, with various frequencies, spatial coverage, and modalities. However, a complete understanding how thalamocortical traveling waves are originated and temporally coordinated in the thalamus and cortex are still unclear. Furthermore, it remains unknown how the network connectivity, excitation/inhibition balance, thalamocortical or corticothalamic delay determine the spatiotemporal wave patterns and characteristics of cortical and thalamic traveling waves. Here we develop a computational model of a two-dimensional thalamocortical network to investigate the thalamic and neocortical traveling wave characteristics in space-time, which allows us to quantitatively assess the impact of thalamocortical network properties on the formation and maintenance of complex traveling wave patterns. Our computational model provides strong theoretical insight into the basis of spatiotemporal wave propagation, as well as experimental predictions that control these wave dynamics.

---

Oscillatory neural activities in the brain are often referred to as brain waves; brain waves propagating across recording electrodes in space are called traveling waves. To date, macroscopic or mesoscopic traveling waves are interpreted as spatiotemporal neural dynamics, which have been reported with various oscillatory frequencies (e.g., theta, alpha, beta, and gamma), spatial coverage (whole brain or local circuits), and modalities (e.g., slice physiology, multielectrode array, high-density EEG or ECoG, and voltage-sensitive dye optical imaging) [1–5]. As a form of either neuronal spiking activity or field potentials, propagating waves have been found in a wide range of cortical, subcortical and thalamic structures, and modulated in a spontaneous or task-dependent manner at different brain states [6–9]. However, it remains challenging to investigate mesoscopic traveling wave patterns based on simultaneous multisite multielectrode-array recordings. It also remains unclear how these spatiotemporal patterns emerge as a result of intra- or inter-network connectivity of local circuits. Biologically-inspired computational models, tightly linked to experimental data, not only provide a complementary approach to investigate these questions, but also offer new experimental predictions on the circuit mechanism of wave propagation [10].

The thalamocortical network and thalamocortical oscillations play important roles in sensory processing, memory consolidation, and multisensory and sensorimotor integration [11, 12]. Rhythmic or synchronous neural activity across various frequency bands has been observed in the thalamocortical system [11]. Propagating wave patterns have also been found in *in vitro* and *in vivo* recordings of thalamic and cortical areas at different brain states [4]. In contrast to continuous and smooth monosynaptic traveling waves observed in the cerebral cortex (CX), traveling waves often appear discontinuous (polysynaptic “lurching wave”) in the thalamus (TH), which involves the reciprocal interaction of thalamocortical (TC) and GABAergic reticular nuclei (RE) cells [13–15]. To generate wave propagation in computational models, traveling waves have been produced in an isolated thalamus or cortex, or in a one-dimensional (1D) thalamocortical system [13, 16]. Nevertheless, the precise nature of how the whole thalamocortical structure, operated as a closed-loop system, determines traveling wave patterns is not completely understood. In addition, it remains puzzling whether the traveling patterns observed in a two-dimensional (2D) topographic network may be preserved in the 1D projection. To date, a wide range of computational modeling work has been developed for traveling waves or spatiotemporal neural activity in neuronal networks [15,17,18], in neural fields [19], and in networks of coupled oscillators [20]. The modeling scale of network size varied between hundreds and tens of thousands of neurons [21]. However, to our best knowledge, no detailed computational model has been yet developed to investigate 2D cortical and thalamic traveling wave patterns and characteristics in a closed-loop thalamocortical system.

Here we develop a biologically-inspired computational model of the 2D topographic thalamocortical network that produces dynamic spatiotemporal patterns from closed-loop interaction of a total of 10,800 cortical and thalamic cell populations. While our proposed 2D network is a reduced version of realistic thalamocortical circuits simplifying several biological details (e.g., Hodgkin-Huxley-type spiking neurons, and laminar cortical structures), it focuses on other important factors, such as the intracortical connectivity, lateral thalamic inhibition, excitation/inhibition (E/I) imbalance, thalamocortical (or corticothalamic) delay, and their impact on the spatiotemporal traveling waves. Our model prediction suggests that rich spatiotemporal patterns and dynamics can emerge independently or jointly from the interactions of these contributing factors. Despite omitting certain biological details, this model demonstrates that spatiotemporal patterns can be interchangeably controlled through alterations in network parameters – without using neural-field approximations that assume a continuous field of neurons. Furthermore, our computational results provide new insight on diseased brain dynamics with regards to abnormality in spatiotemporal patterns.

## RESULTS

### A closed-loop thalamocortical model architecture sustains propagating waves and oscillations

We developed a 2D three-layer thalamocortical network that consists of the cortical and thalamic neurons arranged in a 2D array. For modeling simplicity, we assumed that the cortex collapsed multiple laminar structures into a single-layer structure, with mixture of excitatory (exc) and inhibitory (inh) neurons. The excitatory and inhibitory neurons are connected by excitatory or inhibitory synapses, with certain intracortical connectivity or topography (**Methods**). The thalamus consists of two layers of neurons: excitatory TC neurons and inhibitory RE neurons. Between the thalamus and cortex, there is a bottom-up (feedforward) and a top-down (feedback) connection, forming a closed-loop system (Fig. 1a). There are reciprocal connections between TC and RE. The inhibitory RE→TC projection inhibits the TC neurons; the excitatory TC→RE projection excites RE neurons via axon collaterals, which further inhibits the neighboring TC cells via lateral inhibition. We further ignored RE-RE connections (through dendrodendrite GABEergic synapses), which may disinhibit TC neurons through feedback disinhibition. In the subsequent section, we will explicitly discuss the effect of lateral thalamic inhibition.

**Figure 1.**
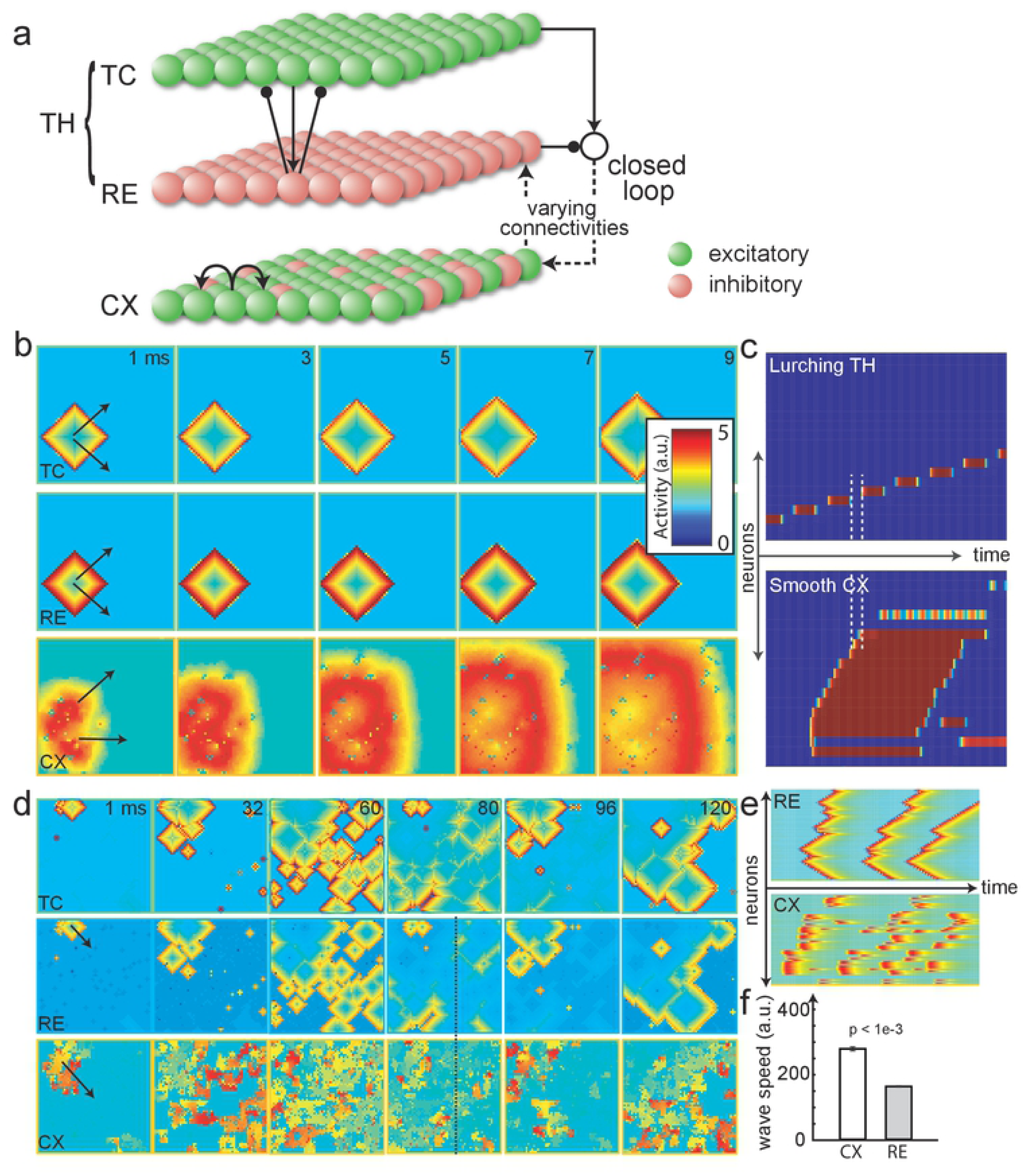
Thalamocortical model and simulated traveling waves. (**a**) Two-dimensional (2D) schematic representation of the computational model with a three-layer architecture: CX (cortex, containing both excitatory and inhibitory cells), RE (inhibitory thalamic reticular nuclei cells) and TC (excitatory thalamocortical relay cells). RE-TC are reciprocally connected. The network is connected in a closed loop. **(b)** Traveling waves produced by the computational model operated in an open loop (i.e., without CX-RE connection, 99% excitatory cortical neurons). Dynamic traveling wave patterns are shown (assuming a dense intracortical connectivity); arrows indicate wave directions. Color bar shows the scale of neuronal activity (a.u.). **(c)** One-dimensional (1D) projection of the traveling waves to indicate the different wave dynamics between the thalamus and the cortex. The gap between the white dashed lines in the thalamus shows the lurching pattern as the wave is staggered in time; in contrast, the cortical wave is smooth. **(d)** Computational model operated in a closed loop (with 10% CX-RE connections and 80% excitatory neurons) results in oscillations with random wave directions. (see **Video S2**) **(e)** 1D projections of CX and RE traveling wave dynamics. (**f**) The average cortical wave speed was significantly faster than the thalamic wave speed (*p*<0.0001 from 5 simulations, student’s t-test). Error bar represents standard error of mean (SEM).

For computational efficiency, we employed an approximation of the FitzHugh-Nagumo (FHN) model and adapted the Izhikevich’s neuron type (Type-1/2/3; see **Methods** and **Fig. S1; Supplementary Table 1**) to simulate the CX/TC/RE neuronal activity at each layer. The TC and RE neurons are of Type-3, and the CX neurons are of Type-1/2. Their model parameters can capture a wide range of input-dependent firing as well as bursting and tonic spiking activity [22].

For the purpose of model validation, we first examined traveling waves in an open-loop condition (i.e., without CX-RE connection, 99% intraconnected CX, 99% excitatory neurons). Our 2D thalamocortical model produced traveling waves in both cortex and thalamus (Fig. 1b). In a 2D graphical illustration, the dynamic evolution of traveling wave was visible in time (from left to right panels, with arrows indicating the wave direction). The mutual recruitment of excitatory spikes and inhibitory rebound spikes created the lurching wave phenotype in the thalamus, resulting in periodic gaps of temporal activity (Fig. 1c), as opposed to the smooth cortical wave, validating the basic structure of our model [4]. The traveling wave initiated at a specific location in the 2D neuronal space, and then spread its pattern across other areas (see 1D projection in Fig. 1c). Next, we examined traveling waves in a closed-loop condition (with RE-CX connection), under a similar connectivity setup. This produced sustained oscillations in both the thalamus and cortex (**Supplementary Video S1**). Note that periodic oscillations were sustained because of the closed-loop feedback and feedforward connections. Without either connection, the thalamic oscillations wouldn’t continue (**Fig. S1d**). Here, the thalamic wave exhibited a diamond-like shape due to the short-range nature of the reciprocal connections. Next, we updated the model conditions to 80% excitatory and 20% inhibitory neurons in the cortex, so as to have the 4:1 ratio of exc-to-inh neurons, and 99% intracortical connectivity. In this case, our model produced oscillations with spontaneous random wave directions (Fig. 1d and **Video S2**). In Figure 1e, the 1D projections illustrate the effect of inhibitory neurons that disrupted the cortical waves through creating inaccessible regions. On average, the cortical wave speed was faster than the thalamic wave speed (Fig. 1f). As the number of excitatory neurons was decreased, the cortical and thalamic wave speeds converged because of the cortical-thalamic synchronization across inhibitory zones (**Fig. S2**). We set the feedback connectivity between the cortex and thalamus at 10%; higher RE-CX connectivity would cause the cortical wave to dominate the thalamic wave, or cause the thalamic lurching (staggered activity in time) to vanish quickly (**Video S3** and **Fig. S2**). Together, these results suggest that by closing the loop, the thalamocortical model may produce rich oscillations with random traveling wave patterns, as well as distinct traveling wave speed between the cortex and thalamus.

### Low intracortical connectivity necessitates clustered cortical neurons to yield traveling waves

In order to sustain wave propagation, or equivalently, to maintain sufficient wave propagation area in time, we found that a high percentage of intracortical connectivity was necessary. In the neocortex, we assumed that the overall intracortical connectivity was 25-36%, with 4:1 ratio of exc-to-inh neurons. In our cortex-alone structure (i.e., without the thalamus) with 25% intracortical connectivity, we found that it was difficult to sustain spontaneous cortical wave propagation. The cortical wave area increased proportionally with the intracortical connectivity (Fig. 2a). When the intracortical connectivity was below 25%, the cortical wave structure lost continuity and reduced to isolated dot patterns. When the thalamus was included in the closed loop, a punctate wave band was observed in the cortex, completely in synchronization with the thalamus (**Fig. S2**), thereby not enabling spontaneous cortical wave propagation.

**Figure 2.**
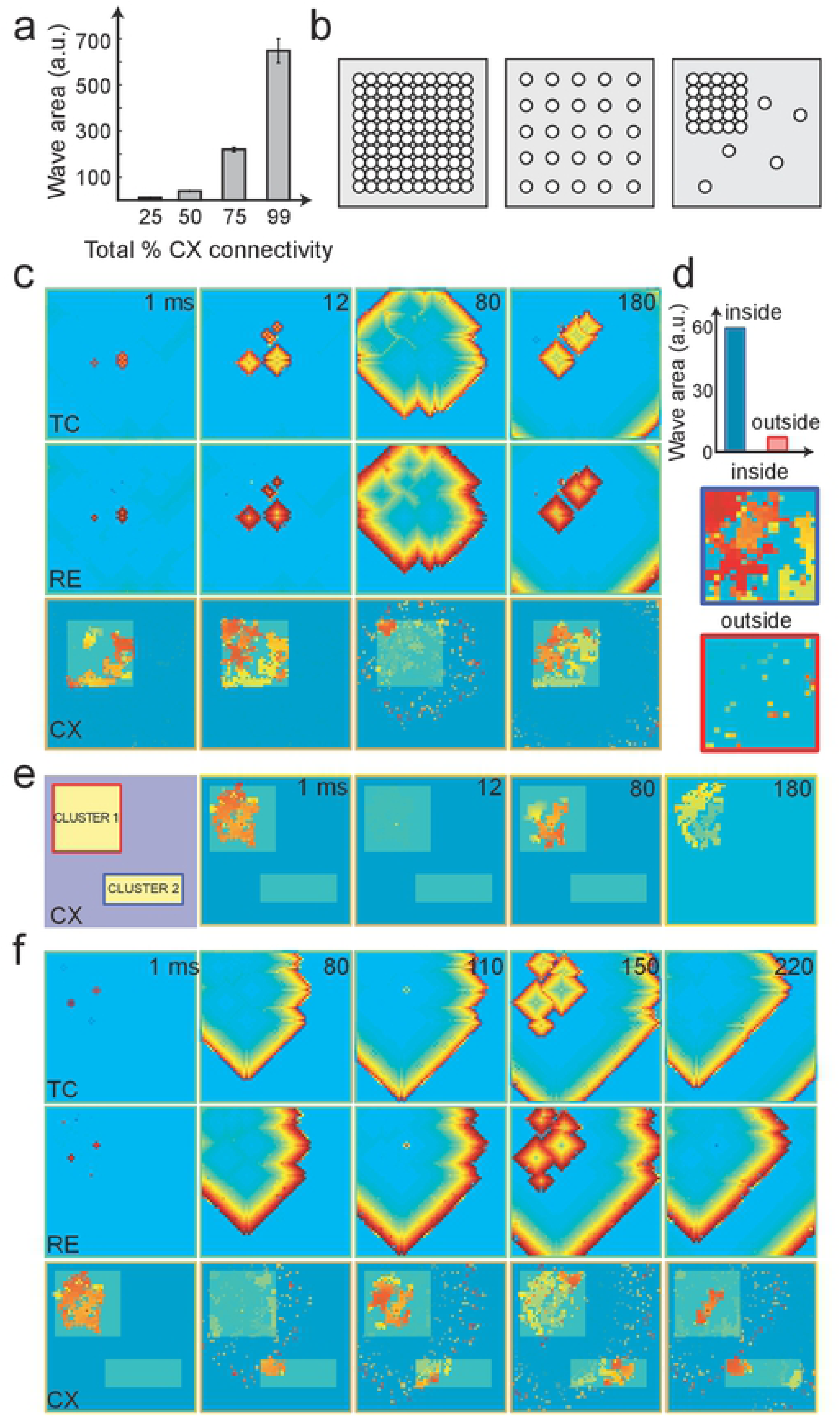
Network connectivity controls patterns of spontaneous traveling waves. (**a)** Traveling wave area was reduced with decreasing overall intracortical connectivity. Error bar represents SEM from 5 simulations. (**b**) Schematic showing different CX arrangement: fully connected (left), uniformly connected with lower connectivity (middle), and clustered and with the same overall connectivity (right). We assumed that the RE and TC layers have uniform arrangement. (**c**) Traveling waves produced by the computational model operated in a closed loop (with overall 25% intracortical connectivity and a 90% intra-connected cluster). The unshaded portion in the CX illustrates the clustered region. (**d)** Comparison of traveling wave area between the inside and outside the clustered region. The wave activity was prominent only within the cluster (zoomed in a snapshot via a blue box), whereas only puncta-type activity was seen outside the cluster (zoomed in a snapshot via a red box). **(e)** Disconnected thalamocortical network with only CX setting: two clusters, both 99% connected, with overall 31% intracortical connectivity. The cluster positions are shown on the leftmost panel. Activity was triggered stochastically within the red cluster. Because of the weak connectivity between two clusters, the activity in the red cluster did not reach the blue cluster. (**f)** Same as panel **e**, except with the thalamus connected in a closed loop. The thalamic wave enabled communications between the two clusters (see **Video S4**).

In cortical circuits, excitatory connections are not uniformly distributed, and exhibit clustering into groups of highly connected neurons [23]. Therefore, for a fixed intracortical connectivity, it implies that the connectivity is high within the clustered groups, and low outside the clustered groups. To investigate the impact of connectivity topography, we modified the 2D arrangement of neurons from uniform connectivity (Fig. 2b, left and middle panels) to clustered structure (Fig. 2b, right panel, with the same overall connectivity as the middle panel). Within the clustered group, the intracortical connectivity was ∼90%, while maintaining the overall 25% connectivity. Our model simulations showed that the propagating waves were prominent within the cluster, as opposed to the puncta patterns outside the cluster (Fig. 2c,d).

Furthermore, we compared the impact of open vs. closed-loop on the cortical traveling waves. In the open-loop condition, we assumed that there was 99% intracortical connectivity within two clustered cortical neuronal groups (with overall 31% connectivity). The cortical wave was initially triggered within cluster 1, but failed to propagate to cluster 2 (Fig. 2e). In the closed-loop setting, under the same connectivity condition, we observed propagating waves in both thalamus and two cortical clusters (Fig. 2f and **Video S4**). This result suggests the potential role of the thalamus in communicating cortical travel waves across multiple isolated areas.

### Lateral thalamic inhibition and thalamocortical delay reshape spatiotemporal cortical dynamics

Next, we considered the RE-RE connections and investigated the effect of lateral thalamic inhibition (within the RE layer). We focused on a reduced thalamocortical model, the RE-CX structure (**Table S2**), where RE cells were fully connected with strong intra-RE excitation, which exerted lateral thalamic inhibition (Fig. 3a). To highlight the effect, we first assumed that CX was fully connected with excitatory neurons only, analogous to the zoom-in view of a clustered group. The RE-CX connectivity was assumed 100% to begin with, and a delay parameter was introduced between RE and CX (i.e., no instant feedback). The time delay during synaptic transmission within a closed-loop system is known to play an important role in its intrinsic dynamics [18, 24].

**Figure 3.**
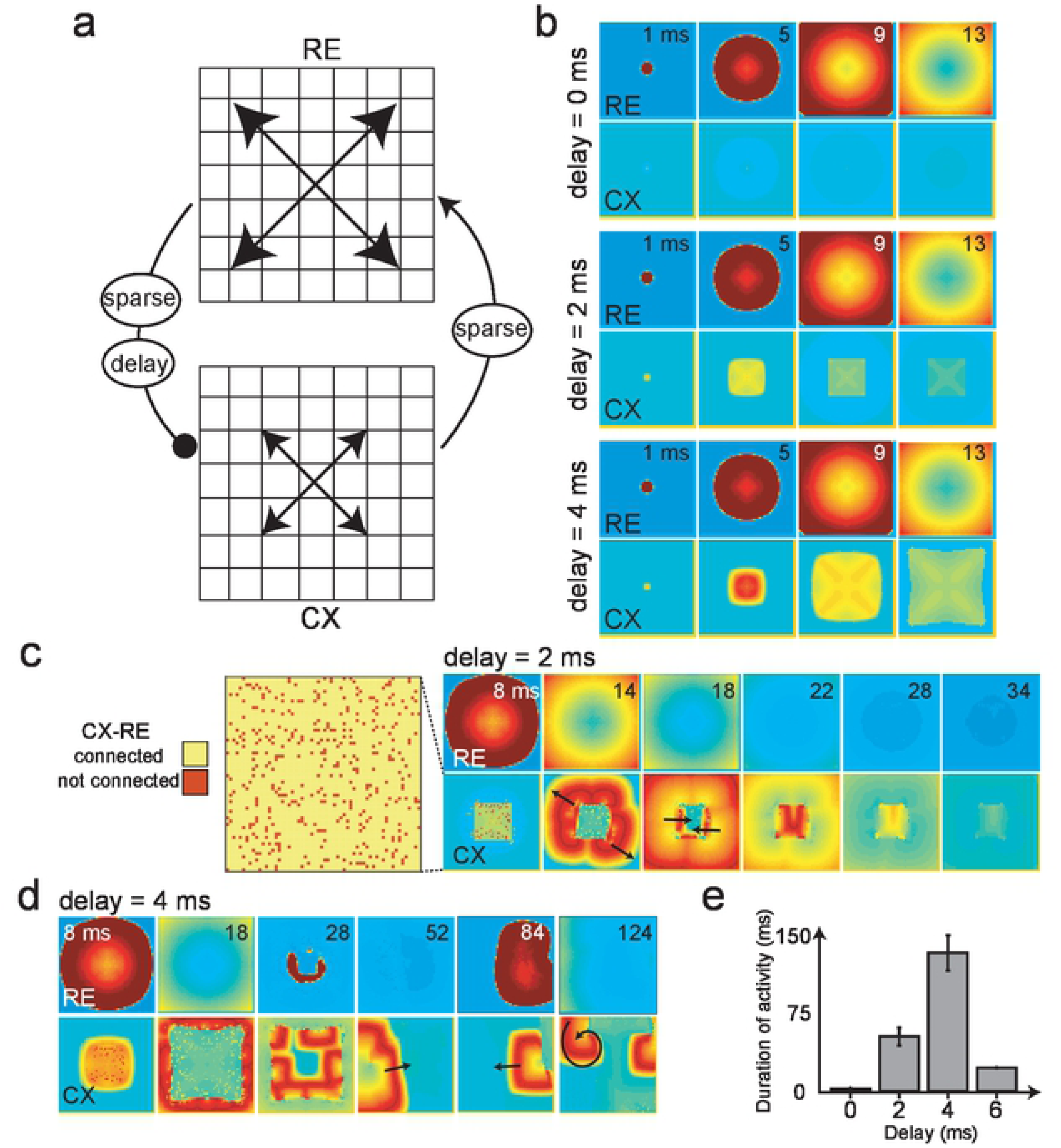
Transmission delay between the cortex and thalamus changes the stimulus-evoked traveling wave patterns. (**a)** A reduced thalamocortical model showing interactions between CX and inhibitory RE cells with lateral inhibition. A nonzero delay parameter was introduced between RE and CX connection to account for axonal conduction delays. We assumed that CX was fully connected with purely excitatory neurons. (**b)** Impact of different thalamocortical delay parameters on CX wave dynamics (assuming fully connected RE-CX). With an increased delay, the CX wave could propagate further and longer. In contrast, lateral inhibition allowed the RE wave to propagate unrestricted regardless the delay. **(c)** A 90% connected RE-CX condition, where the red dots denote the cortical neurons that receive no RE inhibition. For a specific thalamocortical delay of 2 ms, the uninhibited points produced new CX wave that propagated in various directions (indicated by black arrows), and RE wave activity ultimately disappeared (see **Video S5**). (**d)** With an increased delay of 4 ms, dynamic wave activity emerged. In this illustration, radial (t=8 ms), planar (t=52 ms), and rotating (t=124 ms) waves were produced (see **Video S6**). (**e**) Comparison of the wave activity duration with respect to different delay parameters. Their non-monotonic relationship suggests an optimal delay regime in the thalamocortical network. Error bar represents SEM.

We systematically varied the thalamocortical delay parameter (0, 2, 4 ms) and observed the spatiotemporal wave patterns in RE and CX (Fig. 3b). When there was no delay and RE-CX were fully connected, the cortical wave was instantly disrupted by RE inhibition. Increasing the delay to 2 ms allowed the cortical wave to propagate to a certain distance before a complete disruption by RE inhibition. Increasing the delay to 4 ms enabled the cortical wave to propagate further.

In the reduced model, we next lowered the RE-CX connectivity percentage from 100% to 90%, where the unconnected neurons were chosen randomly (Fig. 3c, leftmost panel). In the case of an intermediate delay, we observed a rich repertoire of cortical traveling wave patterns, such as radial, planar, and rotating waves (Fig. 3c,d and **Videos S5 and S6**), and the traveling wave direction or pattern could change in time (t=14 vs. t=18 ms in Fig. 3c; t=52 vs. t=84 vs. t=124 ms in Fig. 3d). The duration of the cortical wave pattern depended on the delay parameter (Fig. 3e). A small delay led to a quick disruption of cortical wave by lateral thalamic inhibition, whereas a large thalamocortical delay caused the cortical wave to escape the field of view before thalamic inhibition became effective. Our results suggest that an optimal delay regime may exist to maintain the cortical traveling wave structure for a thalamocortical network with specific connectivity.

### RE-CX connectivity and thalamocortical delay affect cortical and thalamic wave patterns

In the previous section, we have shown that the randomly-selected unconnected RE-CX nodes produced spontaneous traveling wave patterns. We further investigated whether and how the change of RE-CX connectivity could predict the propagating wave behavior. Using the same setup as in Figure 3b (i.e., nonzero delay and fully connected RE-CX), the cortical wave had a particular range of firing field (dashed box in Fig. 4a) due to RE inhibition.

**Figure 4.**
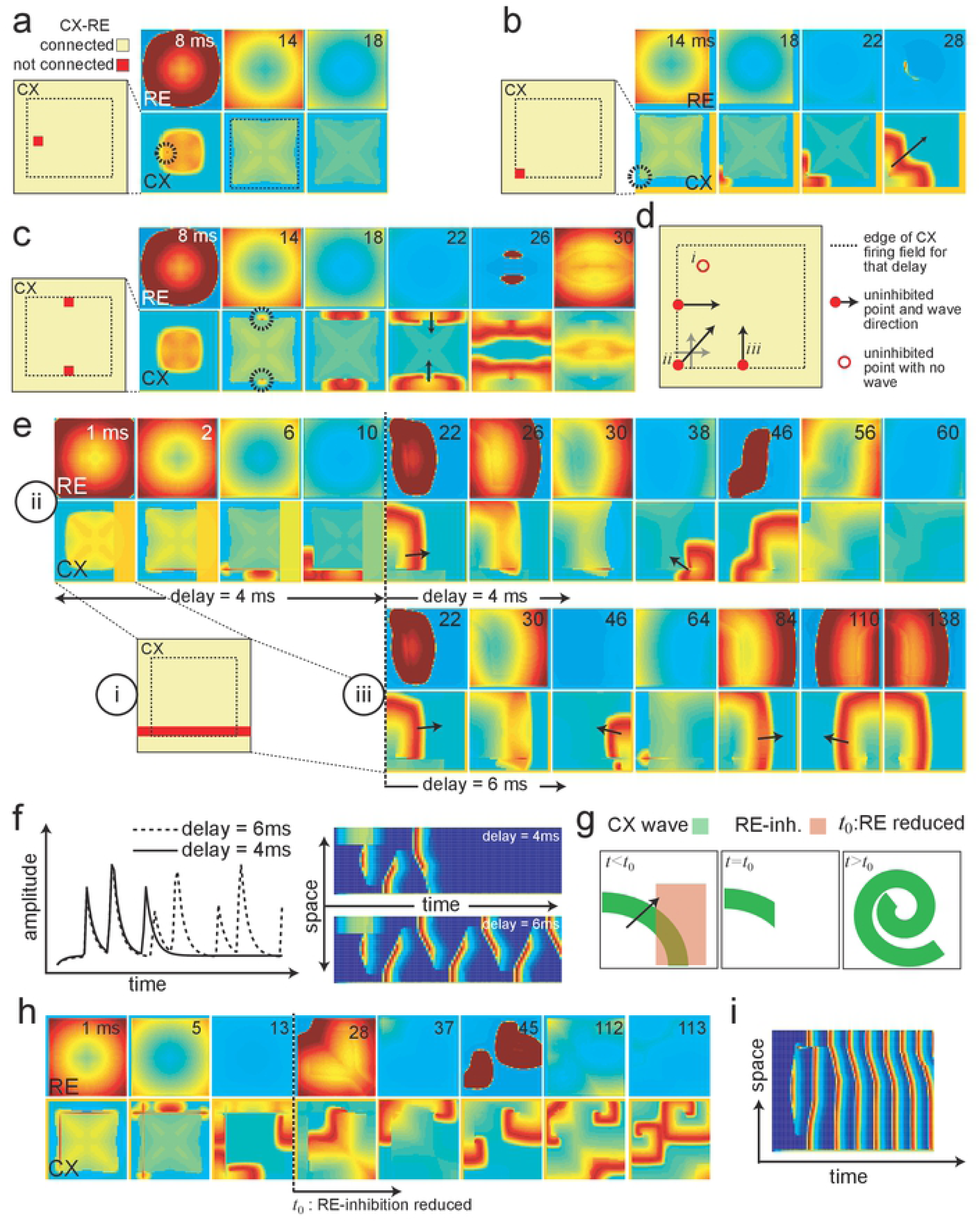
RE-CX connectivity and thalamocortical delays determine cortical and thalamic wave patterns. (**a)** *Left:* A zoomed-in CX circuit showing neurons (red points) that are disconnected to RE. *Right:* Traveling wave dynamics in the CX and RE with lateral RE inhibition. Black circle indicates the uninhibited point. When this uninhibited point fired, it could not produce a wave because of the surrounding CX refractory zone that received RE inhibition. (**b)** Similar as panel **a** except the unconnected point was located at the edge of the CX firing zone (smaller dashed box). Black arrow shows the wave direction. (**c)** Changing unconnected point locations altered wave directions. **(d**) Schematic summarizing how the location of the uninhibited nodes produces traveling waves with various directions. (**e)** *i:* The unconnected points were in a straight line so that the resulting wave oscillated in reverse directions along that line. *ii:* For a delay of 4 ms, two oscillations were allowed until RE disrupted the waves (see **Video S8**). *iii:* By increasing delay (6 ms), infinite oscillations sustained in opposite directions after the initial trigger (see **Video S9**). (**f)** Space-time projections of traveling waves for two delay parameters used in *ii* and *iii*. (**g**) Schematic showing how the RE inhibition can be used to break the CX wave. A broken wave tends to curl around a tip. (**h)** An illustration of generating a rotating cortical wave, which emerged when RE inhibition was reduced at time t_0_ (see **Video S11**). (**i**) Space-time projection of the wave shown in panel **h**.

We considered four distinct scenarios depending on the location and number of the triggered nodes. In the first scenario, in which the unconnected nodes were chosen within the range of cortical firing field, the particular node fired for a longer duration in time because it was not affected by RE inhibition (black dashed circle in Fig. 4a). However, because of the surrounding cortical firing refractoriness, it could not propagate spatially. In the second scenario, in which the unconnected node was at the corner of the cortical firing field (Fig. 4b), a cortical wave emerged in space-time because the cortical firing refractoriness was absent outside the field. The traveling wave initiated outside the dashed box, and then became unconstrained once the refractoriness terminated (**Video S7**). In the third scenario, two unconnected points were selected to create two planar waves in opposing directions. Therefore, for a fixed set of connectivity and delay parameter, the traveling wave patterns (e.g. direction, area and speed) could be predicted (Fig. 4d). On the other hand, when the unconnected node was outside the cortical firing field, no wave was observed. It is important to note that the resulting cortical wave direction was field-size dependent. If there was more room to propagate outside the firing field, then waves could be obtained in other directions as well. The only necessity is that unconnected nodes would need to be at the edge of the cortical firing field.

In the fourth scenario, when the unconnected nodes were chosen in the form of a straight line (Fig. 4e), and the resulting wave oscillated in reverse modes along that line (delay of 4 ms for t=1-10 ms). Under the delay condition of 4 ms, two oscillations could be sustained until the RE inhibition disrupted the cortical wave (t=22-60 ms) (**Video S8**). An initial threshold block was used to initiate a unidirectional wave (Fig. 4e, left panels of CX). If the thalamocortical delay was increased from 4 ms to 6 ms (from t=22 ms on) during the course of simulation (Fig. 4e, bottom), then wave oscillations were sustained indefinitely in opposite directions (**Video S9**). A time-course of these neuronal oscillations are illustrated in Figure 4f, along with a 1D space-time representation. Note that observing these oscillations in time, without a spatial readout, was insufficient to assess the reverse directions.

An interesting observation of traveling wave patterns was the rotating wave produced by the 2D thalamocortical model. Figure 4g illustrates one possible way of producing a rotating wave, in which the RE inhibition was used to break a planar wave, followed by a reduced inhibition level. A broken planar wave has tendency to curl around its tip to create a rotating wave [25–27]. If the RE inhibition was not reduced, the spiral would not get sufficient time to evolve (**Video S10**). Figure 4h shows the implementation of this method where the level of RE inhibition was reduced by ten folds at the moment of t=28 ms. As a consequence of reduced thalamic inhibition, a spiral cortical wave emerged and then continuously repeated itself (**Video S11**). The 1D space-time projection is illustrated in Figure 4i. As seen in our simulations, a 1D space-time representation was insufficient to fully comprehend the underlying spiral wave, and only the 2D traveling wave representation could convey the complete picture. Together, the results suggest that based on the thalamocortical connectivity and transmission delays, one can potentially predict the characteristics of the spatiotemporal patterns that may ensue as a result of perturbations.

### Thalamic and cortical E/I imbalance alters traveling wave frequencies and speeds

The E/I balance in neural circuits is critical for brain functions, and E/I imbalance may induce dysfunctional physiology such as epilepsy and seizures [28]. To investigate the effect of E/I balance on traveling wave characteristics, we focused our attention to a clustered cortical group, in which cortical neurons were nearly fully connected (99% intracortical connectivity).

First, we examined the impact of E/I imbalance in RE inhibition on the cortex. To help illustrate this point, we assumed that the cortex contained 99% excitatory neurons. In a standard closed-loop condition, we showed the 2D thalamic and cortical traveling wave dynamics (Fig. 5a, left), as well as their 1D projections (Fig. 5a, right). The RE and TC competed to trigger the cortical wave activity. In the presence of lower RE inhibition, TC excitation dominated, resulting in firing cortical neurons (Fig. 5a, red dots in white circle). These dots ultimately propagated to form cortical waves (**Video S1**). However, with increased RE inhibition, the effect of TC excitation decreased (due to a lack of firing within the white circle), resulting in fewer cortical traveling waves, or lower frequency (**Video S12**). Comparing the number of striped firing patterns in 1D projections (Fig. 5b vs. Fig. 5a), we observed a decrease in traveling wave frequency induced by increased RE inhibition. When we switched the cortical setup from 99% excitatory neurons back to the 4:1 exc-to-inh neuron ratio, we could still observe a similar effect (Fig. 5c). However, the differences in cortical wave frequency became less prominent as the percentage of excitatory neurons was decreased (Fig. 5d). This result may be ascribed to the fact that inhibitory neurons play a critical role for the oscillatory frequency in the cortex. In contrast, the change in cortical wave speed was insignificant. This insignificant wave speed change was consistent irrespective of the number of excitatory neurons. Adding the effect of lateral RE inhibition to the three-layer model, may potentially alter the wave speed along with frequency.

**Figure 5.**
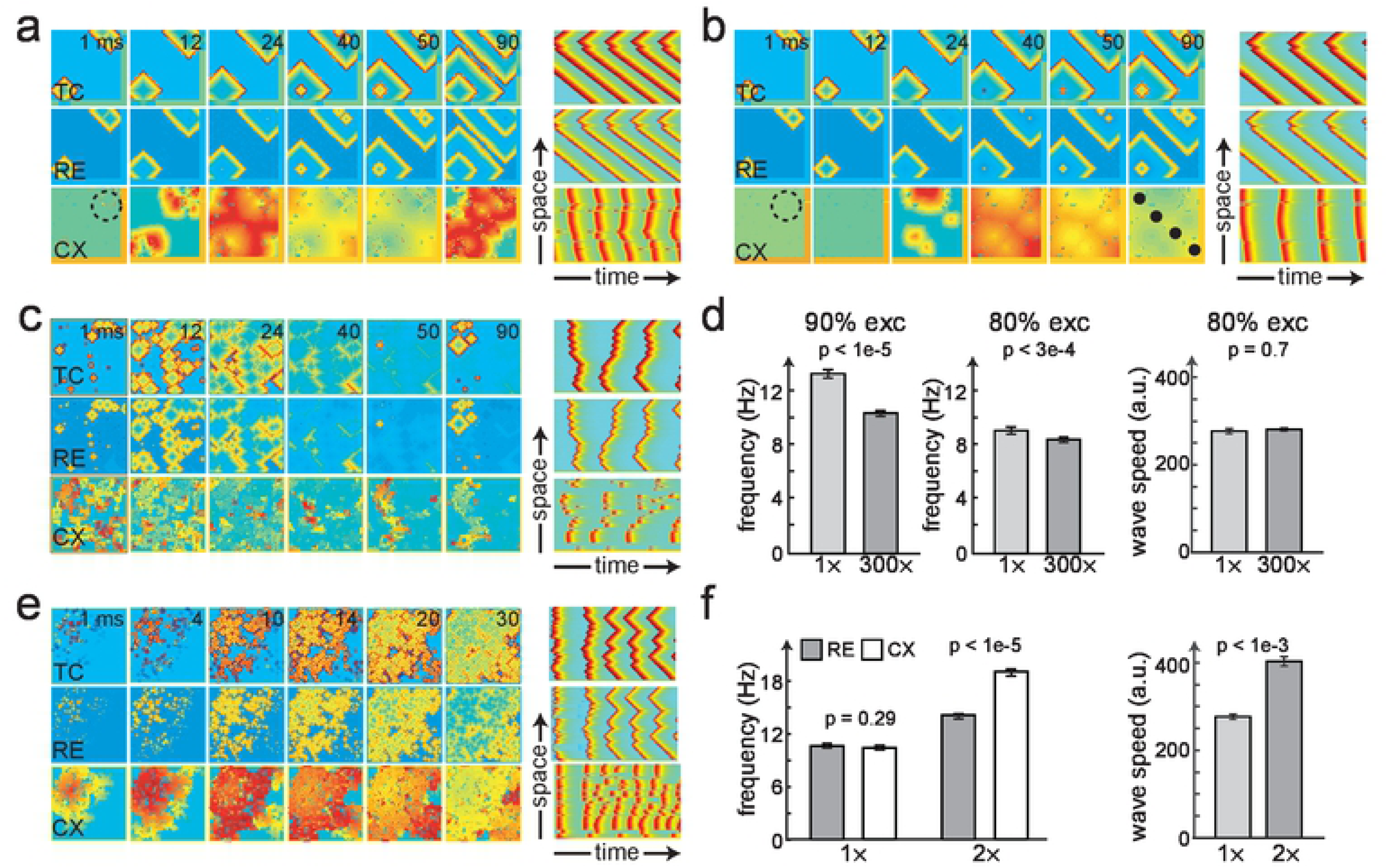
Cortical and thalamic E/I balance alters traveling wave speed and frequency. (**a)** Spatiotemporal activity produced by a closed-loop 2D thalamocortical network (10% RE-CX connectivity, fully connected CX with 100% excitatory neurons). Repeated cortical cell firings occurred due to TC excitatory inputs (dots in the dashed white circle). These dots propagated as a traveling wave (see 1D projection shown on the right). (**b)** RE inhibition weights on CX were increased which reduced the CX firing frequency (compare triggering dots within the white circle in panels **a** and **b**). The black dots in the bottom right panel indicate how spatial frequencies were sampled. (**c)** Similar setup as panel **b** except with 80% cortical excitatory neurons. (**d)** Comparison of the cortical wave speed and oscillation frequency between two levels of RE inhibition (1x vs. 300x). All error bars represent SEM. (**e)** Traveling waves induced by increased CX excitatory weights (10% RE-CX connectivity and 99% intracortical connectivity with 80% excitatory neurons). As seen in space-time projections, wave activity was significantly increased (in comparison with Fig. 1). (**f)** Cortical wave frequency and speed increased as the CX excitatory weights were multiplied by two folds (*p*-values obtained from 5 simulations, student’s t-test).

Next, we examined the effect of imbalance in cortical excitation on traveling waves, by increasing the excitatory weights in the cortex by two folds. As a result, we observed a dramatic change in traveling wave patterns (Fig. 5e**, Video S13**), and a significant increase in both cortical wave frequency and wave speed (Fig. 5f). In the case of 4:1 exc-to-inh neuron ratio and normal model operating conditions, the excitable parameters were assumed such that the thalamus and cortex were synchronized at frequency. However, as we increased the cortical excitation by using a two-fold larger excitatory synaptic weight, the difference between the RE and CX frequencies became prominent (Fig. 5f). Together, the results suggest that E/I balance can alter the traveling wave frequency and speed. Our finding is also in line with the previous 1D model result that the traveling wave speed increases (logarithmically) with the synaptic coupling strength [18].

### Increased cortical variability reduces phase shift in wave activity

Wave propagation is noticeable when there is a phase offset between adjacent firing neurons. This can only occur when a particular set of neurons are triggered and the resulting activity propagates to the neighboring neurons through synaptic connections. This effect will be lost if the nearby neurons also fire spontaneously. This leads us to hypothesize that even when synchronized oscillations are observed, there may be an intrinsic wave pattern that we fail to detect due to the external noise in the network.

To illustrate this point, we conducted computer simulations by calibrating the phase offset between adjacent neurons for varying noise levels at neuronal firing. The sources of external noise could be ascribed to variability in synaptic noise, thermal or conductance noise, and contributions from the modulatory input. We introduced an additional degree of randomness to the basal activity of each neuron in the CX layer (**Methods**). The initiated wave pattern was gleaned from observing the thalamic activity, to which no noise was added. We considered two points in the CX layer (red and blue dots, separated by distance *p*). When the noise level was small (Fig. 6a), the propagating cortical wave was discernible through the time-lapse images, and the phase shift was prominent (Fig. 6a, right panel). Increasing distance *p* led to a greater phase lag. However, when we gradually increased the variance of random noise in the cortical input (Fig. 6b,c), the phase shift decreased or even diminished. Nevertheless, the thalamic traveling wave patterns were still preserved in the latter cases. Together, these results suggest that cortical firing variability would impose challenges to observe the firing phase shift in cortical traveling wave patterns, highlighting the major differences between *in vivo* and *in vitro* conditions.

**Figure 6.**
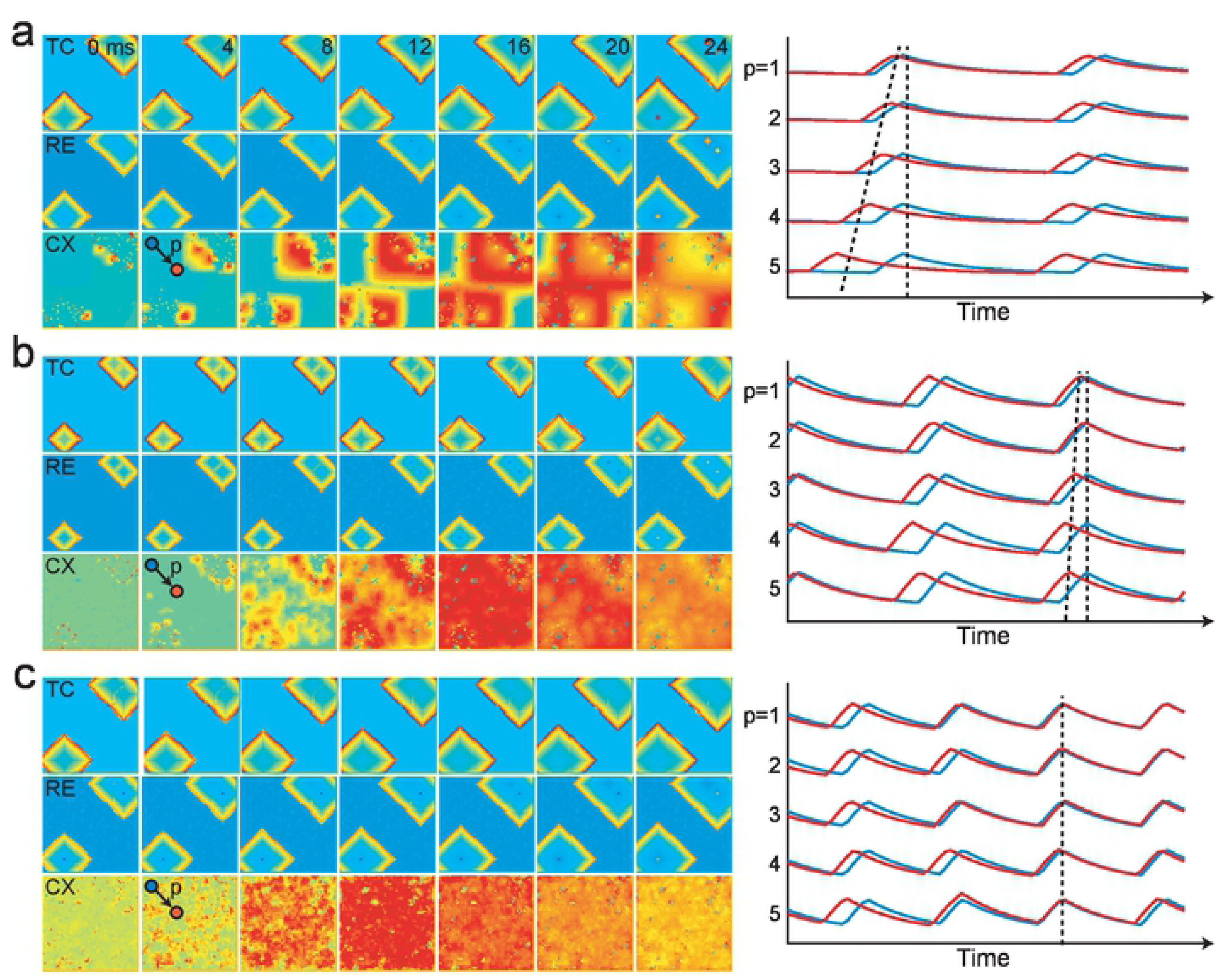
Cortical wave dynamics and phase shift varied with stochastic inputs. (**a)** CX activity (no external noise) was only driven by thalamocortical closed-loop connections initiated with manual triggers, assuming fully connected CX with excitatory neurons. The CX phase shifts in waves were measured by the lag of the activity of the red and blue points. In simulations, the blue point remained unchanged and the red point was moved further away (such that their distance *p* was increased). Five temporal traces (each for a different *p* distance value) of red and blue points are plotted on the right panel. The dashed lines indicate the phase shift as two points move further apart. (**b)** Extra random noise (zero-mean Gaussian with variance 2) was introduced to the CX input (**Methods**). Phase shift was noticeably reduced. (**c)** Additional cortical variability with increased noise variance (from 2 to 4) induced nearly synchronous oscillations regardless of the point location; and the prominent wave structure was completely lost.

## DISCUSSION

We have developed a 2D topographic network of the closed-loop thalamocortical system that produces a broad class of spatiotemporal wave patterns in the cortex and thalamus. Our computer simulations have shown that the propagating wave patterns are influenced by many factors such as intracortical and thalamocortical connectivity, E/I imbalance, thalamic inhibition, and thalamocortical delay. Our results are closely in-line with those generated by the 1D model with delay and spatially decaying connectivity [18]. Furthermore, we show that the 2D traveling waves may display unique characteristics that are indiscernible in 1D projections or *in vitro* (isolated) conditions.

Traveling waves in the brain have been suggested to play important computational and functional roles including memory consolidation, processing of dynamic visual stimuli, sensorimotor integration, and multisensory discrimination and gating [4, 10]. One of speculative roles of macroscopic traveling waves is to propagate and coordinate information across multiple brain regions in space and time; this has been recently verified by a human working memory study using ECoG recordings [29]. Recent experimental findings have shown that thalamic traveling waves may be critical for the development of cortical representations from different sensory modalities [30]. Our results support the hypothesis that a potential role of the thalamus is to enable information being transferred between different cortical areas.

There are two major classes of traveling waves with respect to mechanisms. First, there are spontaneous or internally generated traveling waves, which are generated independent of external stimulus (such as slow waves, spindle waves, and ripple waves) [5,7,8]. The second class consists of stimulus-evoked traveling waves, which are time-locked to the stimulus presented to the sensory or motor system [6,31,32]. While the pattern generation of traveling waves vary, their exact wave patterns or characteristics (speed, direction and area) for information processing in a specific system are still incompletely understood. The speed of cortical traveling waves may have a wide range, ranging from 0.1-0.8 m/s for mesoscopic waves, to 1-10 m/s for macroscopic waves [10, 33]. Multiple factors, including neural recording technique and spatial coverage, may contribute to diverse values of reported traveling wave speed. Our computer simulations predict that the cortical wave speed is influenced by lateral thalamic inhibition and excitatory cortical synaptic connections. In a closed-loop thalamocortical system, increasing the levels of cortical excitation increases the cortical wave speed. Our work also suggests that the inherent degree of noise may conceal the underlying phase shift in the traveling wave activity, yet synchronous oscillations may still retain a wave-nature. This further supports that cortical circuits are functionally two-dimensional [25], and implies the necessity to study 2D traveling wave dynamics or oscillatory spatiotemporal patterns using electrode grid (as opposed to 1D array) recordings.

Transmission delay between adjacent neuronal connections can cause bifurcations resulting in altered dynamics [18, 24]. Our computer simulations have also confirmed that the thalamocortical delay can not only alter spatiotemporal dynamics, but also produce a wide range of traveling wave patterns. The delay parameter elicits a biphasic response, and an optimum delay exists for a particular neural field size that can generate the maximum number and duration of wave patterns. As evidenced from literature [34, 35], the corticothalamic delay is more prominent compared to the thalamocortical delay. Throughout our simulations, we have used a thalamocortical delay to produce traveling wave dynamics. However, as we demonstrated in **Fig. S3**, an asymmetric corticothalamic delay also produced similar cortical traveling wave alterations as the RE-CX connections were changed. In a closed-loop system, the exact location of the delay along the neural pathway (feedforward vs. feedback) did not change the logic behind wave pattern alterations, as the pattern formation theory necessitates only a long-range antagonist that is delayed in time when compared to the local activity [36]. Therefore, with the presence of thalamocortical or corticothalamic delay and the long-range inhibition, a diverse array of traveling wave patterns, including planar, radial, rotating and stationary waves, could be produced from our proposed 2D thalamocortical network. These results suggest that the wave patterns observed in cortical slices may emerge from thalamocortical or intracortical connectivity and altered excitation-inhibition levels rather than purely spontaneous activity.

Previous studies have reported how controlling different parameters of an excitable network can have drastic effects on spatiotemporal patterns. Among the critical parameters are *time-scale separation* (the delay between activator and inhibitor rise-times), *space-scale separation* (faster spreading inhibitor) and *threshold* (excitation-inhibition balance) [27]. Although most studies have considered a uniform excitable medium for altering wave patterns, these parameters can be also altered through manipulating interconnections in a layered system. As we have showed in the current study, the space-scale separation simply amounts to lateral thalamic inhibition that spreads faster than the cortical activity. Similarly, time-scale separation can be equivalent to the corticothalamic delay between long-range connections. The threshold of the system can easily be altered by changing the balance between excitatory and inhibitory synaptic strengths. Therefore, through indirectly manipulating these system parameters, it is theoretically possible to generate a wide range of complex spatiotemporal wave patterns [5, 20].

Through numerical simulations, large-scale computational models may provide insights into the spatiotemporal dynamics of the thalamocortical network at a pathological brain state. The E/I imbalance is an important factor that contributes to epilepsy and seizures [28]. Our results have suggested that in a clustered cortical network, increasing the E/I ratio drastically increases traveling wave speed and overall neuronal excitability, a phenomenon commonly observed in the pathological brain. For instance, traveling waves have been observed during epileptic seizures [37–39], but a complete understanding of their origin remains unclear. One potential mechanism of absence seizure (one kind of primary generalized seizures) is thalamic dysfunction [40–42]. Another plausible mechanism of recurrent seizure is E/I imbalance induced by stronger cortical excitation, which further causes the neuronal network to reach hyperexcitability [43]. Our computer simulations have suggested that the closed-loop thalamocortical system is important for cortical wave propagation, and that the input of excitatory TC cells is necessary to maintain high oscillation frequencies, and subduing TC input through RE inhibition can significantly reduce thalamocortical oscillations. This is consistent with experimental results of a rat model that the thalamus is required to maintain cortical seizure oscillations, and that optogenetic inhibition of TC cell activity disrupts seizure oscillations [44]. Therefore, the dynamic properties of spatiotemporal traveling waves, such as the wave speed, direction and duration, may provide a window to examine pathological brain functions.

Our computational model distinguishes itself from other models in the literature. There are mainly two types of computational mechanisms that produce the traveling waves [20], one is the single neuronal oscillator, and the other based on independent oscillators with their own frequency and phase. Our model is essentially a coupled-oscillator model with short-range synaptic connections, which retains the wave generating characteristics through an all-or-none suprathreshold response. Such a model is capable of sustaining oscillations of different frequencies through Hopf bifurcations, without approximating dynamics through a sinusoidal function as in Kuramoto oscillators. The biophysical details of the Hodgkin-Huxley type neurons are merged into the firing threshold (activation-inhibition balance) of the system, where the network connectivity and synaptic inputs essentially alter neuron’s proximity to the firing threshold. This is a simplifying assumption in order to reduce computation time because numerical simulations of spatiotemporal activity can be computationally cumbersome. Although we have used an approximation of the FHN model in the thalamocortical system, our wave propagation findings should be generalizable to other biophysically-based neuronal models. Many studies in the literature have used approximations of biophysical details in order to analyze spatiotemporal patterns [45], since it is difficult to gain insight from models with high-dimensional parameter space. Because of these simplifications, the exact values of the parameters and their corresponding output wave speed/area values are not as important as is the relative change in the results when system parameters are altered.

In the literature, several 2D computational firing rate models have been developed for cortical structures [25,45–48], but very few have focused on the thalamocortical structure. To date, the available 1D computational models for thalamocortical systems have not explicitly modeled the network connectivity topography (i.e. the clustered intracortical connectivity), or did not jointly model transmission delay and lateral thalamic inhibition [18]. Furthermore, as we show here, the 1D projection has limited characterization capability of traveling wave patterns or properties. A previous 1D computational thalamocortical model has suggested that corticothalamic feedback operates on the thalamus through the excitation of GABAergic RE cells, therefore recruiting TC cells essentially through inhibition and rebound [49]. Although our proposed 2D thalamocortical model keeps this mechanism intact for the thalamus, it does not account for the detailed laminar structure of the cortex, nor does it account for the intra-laminar connections [17,50,51]. It is well known that the superficial layer and deep layer (L5/6) in the cortex have distinct cell density and cell types. The TC cells project to L4, and then propagate the activity to L2/3, and then to L5/6. The corticothalamic feedback initiates from L5/6 and projects back to RE. The corticothalamic collateral input to RE cells is stronger than the direct input to TC cells, emphasizing the modulatory aspect of the feedback. Incorporating more connectivity constraints within the cortical layers would further add details of the modeled neural circuitry [52]. Recent computer simulations have suggested that thalamocortical and intracortical laminar connectivity may change the properties of the sleep spindle wave [51]. Notably, our numerical simulations of 2D thalamocortical network were still on a relatively small scale, and some neurobiological details were not fully modeled. For instance, a separate treatment of thalamocortical and corticothalamic feedback may account for another level of complexity [53]. Furthermore, the CX→TC feedback connection has been omitted since the CX→RE feedback is more predominant and stronger. Also, the bursting behavior of RE cells at rest has been omitted in our current model and only tonic spiking has been focused upon. Introduction of bursting can have interesting effects on the ensuing wave – a topic that needs further exploration. Finally, we have only considered local intra-cellular connections and ignored long-range axonal connections within a layer. These short-range connections, for example, resulted in thalamic waves to be at a 45-degree shape in our numerical simulations. Future work will be required to investigate these issues in greater detail.

## METHODS

### Network architecture and connectivity

We developed a three-layer thalamocortical system by modeling each layer as a 2D sheet of 4-connectivity type (Fig 1a). A layer consisted of a square matrix of neurons. Each neuron had an activator and inhibitor equation (approximation of the FitzHugh-Nagumo or FHN model), along with a synapse output equation. The activator of the model was analogous to the neuron’s membrane potential, while the inhibitor was analogous to the recovery variable. The activator term in the original FHN model is a nonlinear term that allows self-enhancing feedback. We added a saturation approximation to curtail the upper bound of the activator, keeping the nonlinearity intact to allow fast, positive feedback. The inhibitor was the standard FHN-type slow negative feedback, which subdues the activator and creates an ensuing refractory period. The approximation to the FHN model was done to achieve more control over the system output dynamics [54]. Detailed parameters are presented in **Tables S1 and S2.** Although we used an approximation of the FHN model in the thalamocortical system, the wave propagation results are in principle generalizable to other biophysically-based neuronal models.

One feature of thalamocortical network is feedforward inhibition in the cortex. Thalamocortical afferents contact both excitatory neurons and local inhibitory neurons, which synapse on the same excitatory neurons. We assumed that the cortex has 20% inhibitory neurons and 80% excitatory neurons, and has an overall 25% connectivity. The cortex and thalamus are connected in closed loop where a sum of TC and RE activity influences the cortical activity, while the cortex projects feedback to the RE layer. The corticothalamic feedback connectivity between the cortex and the thalamus was set at 10%. Through the corticothalamic feedback, cortical discharges recruited large areas of the thalamus because of the divergent CX-to-RE and RE-to-TC axonal projections. Consequently, the thalamocortical network may generate a wide range of patterns of oscillations and synchrony. The schematic connections within the thalamocortical network are illustrated in **Fig. S4**.

Within the thalamus (TC and RE), there are four types of operations: direct excitation, direct inhibition, lateral inhibition, and feedback disinhibition. Within the cortex, we assumed two types of topological structures (Fig 2b). The first type is a random network with uniform topological structure, and the second type is a locally clustered network.

### Neuron-type and simulation setup

According to neuronal classification of Izhikevich’s criterion [22], we assumed that thalamic neurons belong to Type-3, which generates only a single spike following a step input. This allows neurons to exhibit oscillations when the input is sufficiently large to cause a bifurcation, but the oscillation frequency remains relatively constant for a wide range of input strengths. This neuron also exhibits a rebound spike following an inhibitory synaptic input. The mutual recruitment of excitatory spikes and inhibitory rebound spikes created the lurching wave phenotype that we contrast with the smooth wave [55]. For the purpose of wave propagation, we only considered the tonic firing mode in thalamic neurons. We also assumed that cortical neurons belong to Type-1/2, which exhibits a wide range of oscillation frequencies for different input strengths. This allows cortical neurons to have varying phase oscillations. However, at the bifurcation of Type-1/2 neurons, they do not exhibit rebound spikes to inhibitory synapses.

### **Layer 1: Thalamic relay cells (TC) layer** with activator *T_a_*, inhibitor *T_i_*, synapse *T_s_*

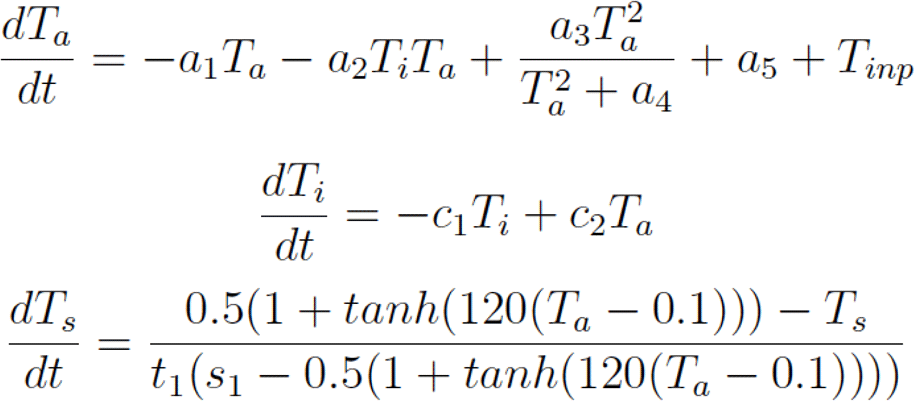

We assumed that the thalamic neurons were set up as Type-3 neuron [22]. The coefficients *a_1_, a_2_, a_3_, a_4_, a_5_* represent the parameters that determine the activator dynamics, more specifically the shape of the activator nullcline (**Fig. S1c**) to give it an inverted-N-shape as is typical of FitzHugh-Nagumo systems. The coefficients *c*_1_ and *c*_2_ determine the slower inhibitor dynamics and produce the inhibitor nullcline a linear shape. The input to the TC layer *T_inp_* consisted of inhibitory synapses from the RE layer (*R_s_*) through a connectivity matrix *C_TH_*.

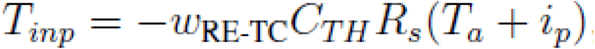

where *C_TH_* connected every TC element to the surrounding four nearest neighboring RE neurons (excluding itself). An inhibitory synapse to TC resulted in a post-inhibitory rebound spike. Each TC neuron sent an excitatory synapse to a corresponding RE neuron, and to the cortical layer.

### Layer 2: Thalamic reticular nuclei (RE) layer with activator *R_a_*, inhibitor *R_i_*, synapse *R_s_*

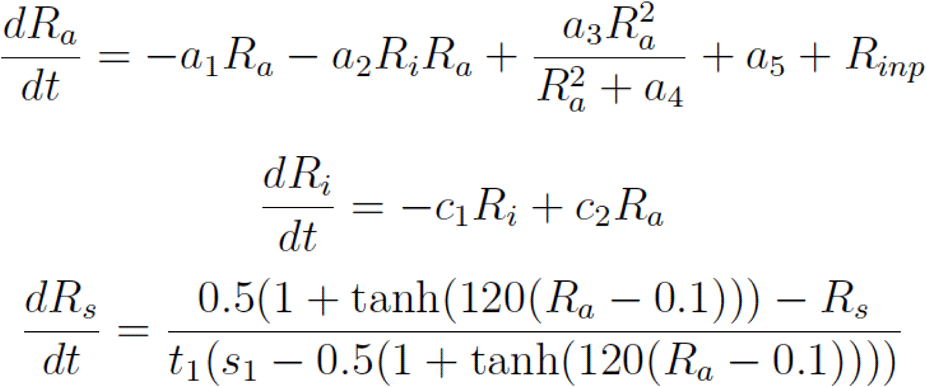

The RE parameters for the activator-inhibitor dynamics were chosen to be same as the TC case for Type-3 neurons. The input to the RE layer *R_inp_* consisted of four parts: (i) excitatory synapses from TC layer; (ii) excitatory corticothalamic inputs from CX; (iii) inhibitory corticothalamic inputs from CX; and (iv) intra-RE excitatory connections that generate lateral thalamic inhibition on the cortex.

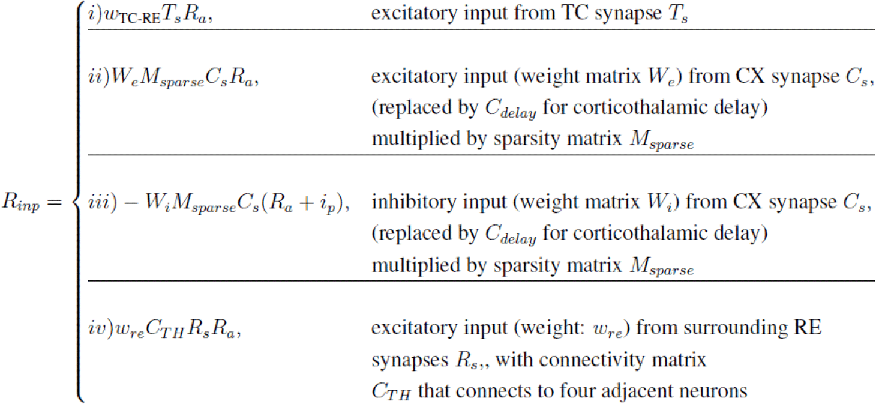

A corticothalamic delay was introduced in the model by using a delayed CX input (*C_delay_* instead of *C_s_*) to the cortex. This was achieved by passing the CX synapse *C_s_* through multiple post-inhibitory rebound spikes before it reached the RE layer. Each delay reaction resulted in a simulation delay of 2 ms. Adding reactions would slow down overall simulation time; therefore, we incorporated a transmission delay up to a maximum of 6 ms. Each delay of 2 ms came from three reactions---exactly similar to those of the Type-3 neurons described in the TC layer with the input being the CX synapse *C_s_*. The delay essentially occurred due to the wait-time before the rebound spike. The delay could be increased by additional 2 ms by incorporating similar rebound-spike delays, using the first delay synapse as the input, and so on.

The weight matrices *W_e_* and *W_i_* consisted of the excitatory and inhibitory weights from the excitatory and inhibitory neurons in the cortex, respectively. We assumed that the RE-CX connections were symmetric and sparse, as characterized by a binary matrix *M_sparse_*. The cortical input to the TC layer was considered negligible when compared to the corticothalamic input to RE. The fourth input, intra-RE excitation, was used only in the reduced RE-CX model case (when the input (i) in *R_inp_* was removed), which was aimed to mimic the effect of lateral RE inhibition on CX. Furthermore, the net input to a RE neuron was saturated at a minimum to prevent numerical instabilities.

### **Layer 3: Cortex (CX) layer** with activator *C_a_*, inhibitor *C_i_*, synapse *C_s_*

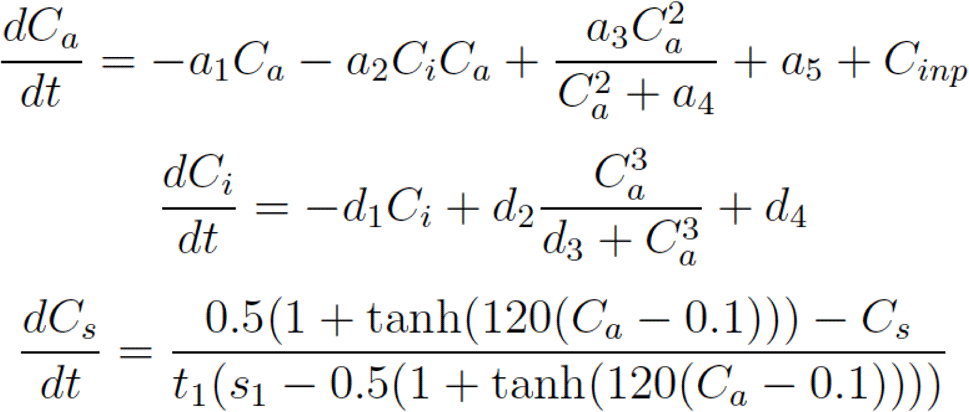

The cortical inhibitory neurons were set up as Type-1/2 neuron, which do not exhibit the inhibitory postsynaptic rebound. Instead, they incorporate saddle-node bifurcations that allow different spiking frequencies for different input strengths. The input to the CX layer *C_inp_* consisted of five parts: (i) excitatory synapses from TC layer; (ii) inhibitory inputs from RE layer with a delay parameter; (ii) excitatory inputs from neighboring cortex neurons; (iv) inhibitory inputs from neighboring cortex neurons; and (v) random Gaussian noise input.

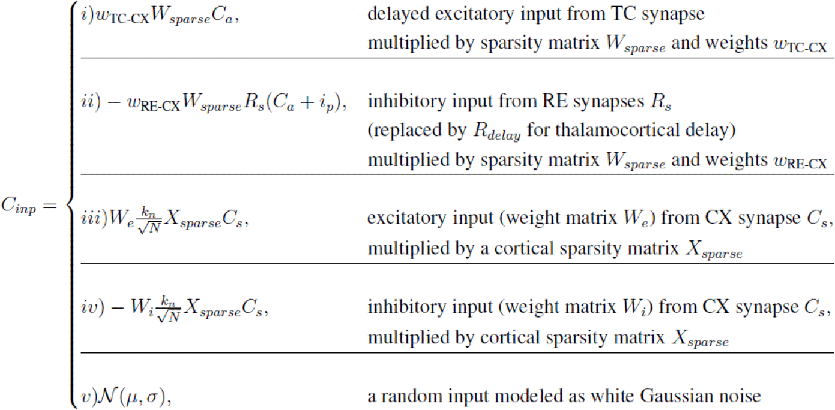

A thalamocortical delay was introduced in the model by using a delayed RE input (*R_delay_* instead of *R_s_*) to the cortex. This was achieved by passing the RE synapse *R_s_* through multiple post-inhibitory rebound spikes before it reached the cortex. Each delay reaction resulted in a simulation delay of 2 ms. Adding reactions slowed down overall simulation time, therefore, we incorporated a transmission delay up to a maximum of 6 ms. Each delay of 2 ms came from three reactions---exactly similar to those of the Type-3 neurons described in the TC layer with the input being the RE synapse *R_s_*. The delay occurred due to the wait-time before the rebound spike. The delay could be increased by additional 2 ms by incorporating similar rebound-spike delays, using the first delay synapse as the input, and so on.

We assumed that the intracortical connectivity was sparse (characterized by a binary and symmetric matrix *X_sparse_*), which incorporated the effect of clustering or rearrangement of cortical cell connections. For modeling simplicity, we assumed that the thalamocortical connections were symmetric (i.e., *W_sparse_* = *M_sparse_*). We also assumed that the exc-to-inh neuron ratio was 4:1 in the cortex.

### Numerical simulations and traveling speed characterization

The neural equations were numerically simulated using the SDE toolbox of MATLAB (MathWorks) [56], using a 2D array for a neural layer with absorbing (high-threshold) boundary conditions. Kymographs were calculated from the 2D activity through line-scans in time. In our model, each layer consisted of 3,600 neurons (array of 60×60), the computer simulation time for the proposed three-layer network was ∼600 s for 400-ms duration. Statistics were done by varying the (feedback, feedforward and intracortical) network connectivity and the arrangement of excitatory neurons in the cortex. Student’s t-test was used to calculate statistical significance.

To measure the traveling wave speed, a custom MATLAB script was used. Briefly, the wave front was segmented at subsequent frames of videos. Based on these segmentations, the number of patches and averaged area was computed. To measure the averaged wave speed, the distance from each pixel on the boundary of a wave front in frame *n*+1 to the closest edge of a wave in frame *n* was computed. Our computed wave speed was in arbitrary unit (a.u.). To put that in a perspective, if we record 60×60 array neuronal activity from a 2D multielectrode array (MEA) of ∼24×24 mm^2^, then each frame of the wave videos reflects the 2-ms temporal resolution such that the simulated wave speed is ∼100-300 cm/s.

## ACKNOWLEDGMENTS

We thank M. Bazhenov, J.D. Murray, Y. Hu, J. Rinzel, and W. Truccolo for their valuable comments. This research was partially supported by the DAPRA under contract number HR0011-16-0139 (P.A.I.), NSF grant CBET-1835000 (Z.S.C.), NIH grants R01-NS100065 (Z.S.C.) and R01-MH118928 (Z.S.C.). The funders had no role in study design, data collection and analysis, decision to publish, or preparation of the manuscript.

## AUTHOR CONTRIBUTIONS

P.A.I. and Z.S.C. conceived the study. The model was developed by S.B and P.A.I aided by M.B.L.C. S.B. carried out the simulations. S.B., P.A.I and Z.S.C. wrote the manuscript.

## Supporting Information

**Fig S1.** (**a**) Diagram of the thalamocortical circuit (adapted from Destexhe and Contreras, 2011). (**b**) Illustration of the open and closed-loop connections between the CX, TC and RE. (**c**) Illustrations of phase plane for Type-1/2 and Type-3 neurons (Izhikevich’s classification) used in our computer simulations. **(d)** Comparison of open-loop (left) and closed-loop (right) traveling wave dynamics (shown in the form of space-time projections) of the thalamocortical system when both the thalamus and cortex were manually triggered at a point initially.

**Fig S2.** (**a**) Illustration of how lurching behavior of thalamus was lost with different RE-CX connectivity percentages (10%, 50% and 90%). A higher percentage of CX-RE connectivity eliminated the time-gap between subsequent triggers (highlighted in white dashed circle) due to cortical inputs. (**b**) Illustration of the CX and RE traveling waves with overall 25% intracortical connectivity. In this case, the cortical wave was punctate and discontinuous, thereby being difficult to detect. (**c**) As the percentage of excitatory cortical neurons was reduced, the CX wave speed decreased, whereas the RE wave speed increased slightly. Error bars represent SEM (n=5 simulations).

**Fig S3. (a)** The RE-CX model with corticothalamic delay (in comparison with the thalamocortical delay shown in Figure 3a). **(b)** Effect of different corticothalamic delay parameters on cortical wave patterns. **(c)** An instance of cortical traveling wave dynamics using a corticothalamic delay of 2 ms. **(d)** Illustration of diverse cortical and thalamic wave pattern formation using a sparse random connectivity matrix. **(e)** An instance of generating rotating cortical wave (similar to Figure 4h). RE inhibition was reduced at time t_0_.

**Fig S4.** Graphical illustration of thalamocortical network connections, with green indicating excitatory (exc), red indicating inhibitory (inh), and blue indicating mixed excitatory and inhibitory connections, respectively. *T_s_, R_s_*, and *C_s_* represent the synapses for the TC, RE and CX layers, respectively. The number inside the circle represents the connectivity percentage.

## Supplementary Video Legend

**Video S1**: Closed-loop spontaneous oscillations of the three-layer thalamocortical system with 60×60 neurons in each layer (99% intracortical connectivity, 99% excitatory neurons, 10% Cortex-Thalamus connections). Wave directions are indicated by the vector field using the optical flow method in MATLAB.

**Video S2**: Closed-loop spontaneous oscillations of the three-layer thalamocortical system with 60×60 neurons in each layer (99% intracortical connectivity, 80% excitatory neurons, 10% Cortex-Thalamus connections).

**Video S3**: Closed-loop spontaneous oscillations of the three-layer thalamocortical system with 60×60 neurons in each layer (99% intracortical connectivity, 80% excitatory neurons, 90% Cortex-Thalamus connections).

**Video S4:** Closed loop oscillations with two highly connected clusters in the cortex (80% excitatory neurons, 10% Cortex-Thalamus connections).

**Video S5:** Reduced model CX-RE. Dynamics with 90% CX-RE connections and a delay of 2 ms.

**Video S6:** Reduced model CX-RE. Dynamics with 90% CX-RE connections and a delay of 4 ms.

**Video S7**: Reduced model CX-RE. Dynamics with one unconnected point of CX-RE (24 ms), which creates a traveling wave in the diagonal direction. Delay of 4 ms.

**Video S8**: Reduced model CX-RE. Dynamics with a line of unconnected CX-RE points (24 ms), that creates oscillating planar waves. The initial threshold block is to create a unidirectional planar wave (t<24 ms). Delay of 4 ms.

**Video S9:** Reduced model CX-RE. Dynamics with a line of unconnected CX-RE points (24 ms), that creates oscillating planar waves that are sustained infinitely. Delay is increased from 4 ms to 6 ms at t=40 ms.

**Video S10:** Reduced model CX-RE. Dynamics with two lines of unconnected CX-RE points (24 ms), which are used to break the planar wave. The wave starts to rotate before being subdued by RE inhibition. Delay of 4 ms.

**Video S11:** Reduced model CX-RE. Dynamics with two lines of unconnected CX-RE points (24 ms), which are used to break the planar wave. The wave starts to rotate and ultimately evolves into two rotating spirals as the RE inhibition is reduced after t=48 ms. Delay of 4 ms.

**Video S12**: Closed-loop spontaneous oscillations of the three-layer thalamocortical system with 60×60 neurons in each layer (99% intracortical connectivity, 99% excitatory neurons, 10% Cortex-Thalamus connections), with RE inhibition increased.

**Video S13**: Closed-loop spontaneous oscillations of the three-layer thalamocortical system with 60×60 neurons in each layer (99% intracortical connectivity, 80% excitatory neurons, 10% Cortex-Thalamus connections), with intracortical excitatory weights increased.

